# Multimodal Pretraining for Unsupervised Protein Representation Learning

**DOI:** 10.1101/2023.11.29.569288

**Authors:** Viet Thanh Duy Nguyen, Truong Son Hy

## Abstract

In this paper, we introduce a framework of symmetry-preserving multimodal pretraining to learn a unified representation of proteins in an unsupervised manner, encompassing both primary and tertiary structures. Our approach involves proposing specific pretraining methods for sequences, graphs, and 3D point clouds associated with each protein structure, leveraging the power of large language models and generative models. We present a novel way to combining representations from multiple sources of information into a single global representation for proteins. We carefully analyze the performance of our framework in the pretraining tasks. For the fine-tuning tasks, our experiments have shown that our new multimodal representation can achieve competitive results in protein-ligand binding affinity prediction, protein fold classification, enzyme identification and mutation stability prediction. We expect that this work will accelerate future research in proteins. Our source code in PyTorch deep learning framework is publicly available at https://github.com/HySonLab/Protein_Pretrain.

## INTRODUCTION

Proteins, the essential building blocks of life, play a crucial role in a wide range of biological processes, rendering them a subject of profound scientific interest. Understanding the intricate structures and functions of proteins holds immense significance, yielding valuable contributions to numerous fields, such as molecular biology, medical research and drug design [1, 2]. The advent of data-driven approaches, including machine learning (ML) and deep learning (DL), has revolutionized the field of protein research [3, 4]. These methods leverage data to unlock deeper insights into proteins, offering more precise predictions while significantly reducing the need for resource-intensive laboratory experiments.

While supervised representation learning in protein research has made considerable progress, its potential remains constrained by the limited availability of labeled data, a resource-intensive and time-consuming requirement. As a result, there is a growing interest in unsupervised pretraining methods, which can equip models with foundational knowledge of proteins without relying on extensive labeled datasets. Inspired by the remarkable success of unsupervised pretraining in domains like Natural Language Processing (NLP) [5] and generative AI [6], researchers have increasingly turned their attention to applying similar techniques to proteins, aiming to learn representations that capture both their structural intricacies and functional characteristics. These efforts have yielded notable achievements in advancing our understanding of proteins [7, 8].

Proteins are complex biomolecules with a hierarchical structure, consisting of four distinct levels: primary, secondary, tertiary, and quaternary (as illustrated in Figure 1). Each of these structural levels corresponds to a specific modality of representation. While previous pretraining methods have typically treated these modalities in isolation, the complexity and multifaceted nature of proteins necessitate a more comprehensive and integrated approach to representation learning. Furthermore, these methods have commonly disregarded the critical aspect of preserving symmetries inherent to proteins, including permutation, rotation, and translation. Proteins exhibit symmetrical properties crucial to their biological functions, and failing to account for and maintain these symmetries can lead to inaccuracies in representation. To address the limitations of previous pretraining methods, we propose dedicated unsupervised symmetry-preserving pretraining methods for each modality of protein. Specifically, for amino acid sequences, we use Evolutionary Scale Modeling (ESM-2) [9], a state-of-the-art method for learning representations of unaligned protein sequences that take into account evolutionary information; for residue-level graphs and 3D point clouds, we use Variational Graph Auto-Encoders (VGAE) [10] and PointNet Autoencoder (PAE) [11], respectively, to learn representations that can capture the spatial relationships between residues and atoms. These methods are all designed to capture the unique characteristics of each protein modality, while also preserving the symmetries that are inherent to proteins. Additionally, we utilize Auto-Fusion [12] to synthesize a joint representation from these pretrained models, encouraging effective intermodal information extraction. All of our contributions allow us to produce more informative, robust, and unified representation of proteins, which can lead to significant improvements in a variety of protein-related tasks.

**FIG. 1.**
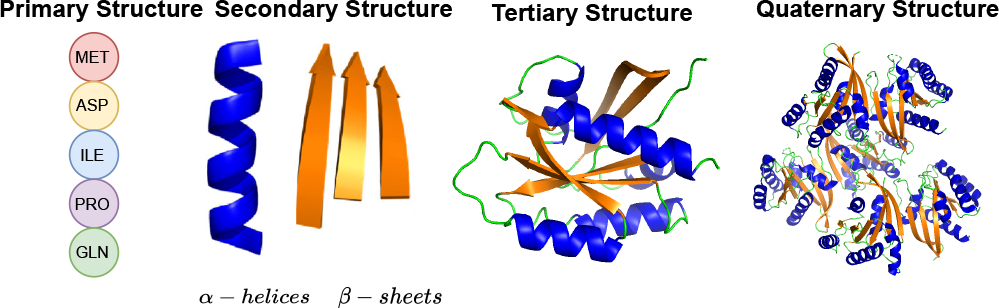
Different levels of protein structure including primary, secondary, tertiary, and quaternary structure. At the primary structure, a protein is simply a chain of amino acids. The secondary structure refers to the local folding of the protein chain into alpha helices and beta sheets. The tertiary structure refers to the overall 3D structure of the protein, which is determined by the interactions between the different amino acids in the protein chain. The quaternary structure refers to the interactions between multiple protein subunits to form larger protein complexes.

Our contributions can be summarized as follows:

- Developing dedicated unsupervised symmetry-preserving pretraining methods for each protein modality, specifically applying Evolutionary Scale Modeling (ESM-2) to amino acid sequences, Variational Graph Auto-Encoders (VGAE) to residue-level graphs, and PointNet Autoencoder (PAE) to 3D point clouds.
- Leveraging Auto-Fusion to synthesize joint representations from these pretrained models, which encourages effective intermodal information extraction.
- Carefully analyzing the performance of our framework in the pretraining tasks to provide a comprehensive evaluation.
- Demonstrating the effectiveness of our pretrained protein representation on several downstream tasks including protein-ligand binding affinity prediction, protein fold classification, enzyme identification and mutation stability prediction.

## II. RELATED WORK

### A. Supervised representation learning on proteins

Supervised representation learning on proteins aims to develop discriminative models by training them on labeled datasets tailored for specific protein-related tasks, such as predicting protein binding interfaces [13], deciphering protein functions [14], or estimating protein-ligand binding affinities [15]. These models are designed to make predictions or classifications based on the input data, effectively mapping protein-related information to predefined labels.

While supervised methods have shown success in these specialized applications, they have notable limitations. One significant limitation is that these models tend to learn task-specific representations. This means that the learned representations are often fine-tuned for a particular task and may not generalize well to other protein-related tasks or provide a broader understanding of proteins. They lack the capacity to capture the diverse and intricate aspects of protein structures and functions that go beyond the specific tasks for which they were trained. Another limitation of supervised learning methods is their substantial data requirements. These models demand extensive labeled data for effective training, which can be particularly challenging in the domain of protein-related tasks. Collecting large amounts of labeled protein data is not only costly but also time-consuming, making it a significant obstacle to the widespread application of supervised methods in this field. In contrast, unsupervised pretraining methods can address the limitations of supervised learning methods by learning representations of proteins from unlabeled data, which is more abundant and easier to obtain than labeled data. Additionally, it can learn more general-purpose representations of proteins that can be used for a variety of tasks.

### B. Pretraining on sequences

Proteins are composed of long chains of amino acids, known as the amino acid sequence, which serve as foundational information about these biological macromolecules. Learning from amino acid sequences is critical because they encode the intricate blueprint for protein structure and function. The sequence determines the folding of the protein and governs its interactions with other molecules, playing a pivotal role in the exploration of molecular biology. Drawing inspiration from the remarkable success of language models in the field of Natural Language Processing (NLP), the computational biology community has ventured into developing protein language models [8]. Much like language models, which encode semantic meaning from text, protein language models aim to extract semantically rich representations from protein sequences. All Protein Language Models (PLMs) are pretrained on large datasets of protein sequences through unsupervised tasks, which allow the models to learn rich representations of protein sequences and their underlying features.

Several iterations of PLMs have been developed in recent years, with ESM-1 [16], the Evolutionary-Scale Modeling, standing out as a notable example. ESM-1 has been extensively trained on an impressive dataset comprising 250 million protein sequences, which amounts to a staggering 86 billion amino acids. The training process involved masked-language-modelling tasks, a prevalent method among PLMs. Additionally, ESM-2 [9], a more recent development building upon the ESM-1 architecture, offers variations with parameter counts ranging from 8 million to a staggering 15 billion. Another noteworthy PLM is ProtTrans T5 [17], drawing inspiration from the T5 model [18] originally designed for natural language processing. This PLM takes the form of a 3 billion parameter encoder-decoder model and was trained with a denoising task where 15% of the amino acids in the input were randomly masked. Furthermore, Protein-BERT [19] is inspired by BERT, but contains several innovations such as global-attention layers that have linear complexity for sequence length. As a result, the model can process protein sequences of almost any length, including extremely long protein sequences. These models all share a common objective in their training process, focusing on masked-token prediction. In addition, there is an alternative pre-training approach in the realm of protein language models that focuses on contrastive learning. Notably, CPCProt [20] employs the principle of mutual information maximization between local and global information as a self-supervised pretraining signal for protein embeddings. This approach involves dividing protein sequences into fixed size fragments and training an autoregressive model to distinguish between subsequent fragments from the same protein and fragments from random proteins.

While PLMs have achieved impressive results on a variety of downstream tasks for structure and function prediction [21, 22], they are bound by the inherent limitations of amino acid sequence data alone. Amino acid sequences can reveal primary structural information, including the order and type of amino acids in a protein chain. However, they fall short in depicting the intricate interactions between amino acids, the spatial arrangements of atoms within a protein, and the dynamic three-dimensional structures that proteins adopt. These limitations extend to the inability to elucidate how proteins interact with ligands, cofactors, and other biomolecules, which is crucial for understanding their functions.

### C. Pretraining on residue-level graphs

Graph neural networks (GNNs) utilizing various ways of generalizing the concept of convolution to graphs have been widely applied to many learning tasks, including modeling physical systems [23], finding molecular representations to estimate quantum chemical computation [24–27], etc. One of the most popular types of GNNs is message passing neural networks (MPNNs) [28] that are constructed based on the message passing scheme in which each node exchanges information to its neighborhood. In our work, we utilize existing deep generative models on graphs, in particular variants of graph variational autoencoder [10, 29], to pretrain on the residue-level graphs of proteins.

Proteins are intricate three-dimensional structures where the spatial proximity of amino acids (residues) holds great significance. Relying solely on amino acid sequences for learning may lead to an oversight of these spatial relationships, thereby limiting the efficacy of sequence-based methods. It is important to recognize that two residues, despite being distant along the protein sequence, could actually be positioned closely in three-dimensional space. This discrepancy arises because protein sequences undergo complex folding processes, and the linear order of amino acids in the sequence does not always mirror their relative positions in the three-dimensional structure. Pretraining on residue-level graphs (i.e. K-nearest-neighbor graph in which each node is a residue and the edge is determined by the Euclidean distance between two corresponding residues), which effectively capture the interconnections between residues, provides researchers with the means to gain essential insights into the structural and functional aspects of proteins. This approach enriches our comprehension of protein structures and empowers us to model proteins more effectively.

Several research initiatives have delved into the application of residue-level graphs in the domain of protein representation learning. Notably, Multiresolution Graph Transformers (MGT) [30] introduced the pioneering concept of a graph transformer architecture capable of learning representations for large molecules at multiple scales, encompassing proteins, peptides, and polymers. MGT excels in learning atom-level representations, allowing for the meaningful grouping of atoms into functional units. Another innovative development, ProNet [31], is a hierarchical graph network designed to elevate protein representation learning, with a specific emphasis on 3D structures. ProNet enables the computation of protein representations at varying levels of granularity, with each amino acid serving as a node within the graph model. Additionally, GearNet [32] was proposed as an elegant yet highly effective structure-based encoder for protein representation learning. This approach relies on relational message passing over protein residue graphs and harnesses self-supervised pretraining techniques, including multiview contrastive learning and self-prediction tasks.

The utility of residue-level graphs in protein representation learning is under exploration, and significant results have been achieved in a variety of protein-related tasks. However, pretraining on residue-level graphs is not without limitations. While significant advantages in capturing spatial information about protein structures are offered by these graphs, their ability to capture long-range interactions in large protein structures is limited due to their reliance on local interactions. Effectiveness in representing local structural patterns and interactions between nearby residues is achieved by graph neural networks built based on the message passing scheme, but they may struggle to capture the global structural context or long-range dependencies in proteins [33, 34].

### D. Pretraining on 3D point clouds

While amino acid sequences and residue-level graphs provide valuable information about proteins, they have inherent limitations in capturing the detailed spatial arrangements of atoms and the intricate folding patterns that define a protein’s shape. Recent advancements in deep learning have introduced unsupervised representation learning techniques that leverage 3D point cloud data [35], a modality that is relatively unexplored in the context of protein representation learning. This novel approach harnesses the power of 3D point clouds to gain insights into the precise positions of atoms within proteins and their dynamic structural changes.

In the field of computer vision, the utilization of 3D point clouds for pretraining has led to remarkable advancements. It has not only transformed tasks such as object recognition, scene understanding, and autonomous navigation but has also underscored its potential for capturing and comprehending intricate three-dimensional environments. Notably, an innovative self-supervised pretraining method, as presented in [36], operates seamlessly with single-view depth scans obtained from a variety of sensors, eliminating the need for 3D registration and point correspondences. This approach efficiently pretrains standard point cloud and voxel-based model architectures. Conversely, the introduction of Point-M2AE [37], a robust Multi-scale Masked Auto-Encoder (MAE) pretraining framework designed for hierarchical self-supervised learning of 3D point clouds, offers an effective means of capturing spatial geometries and semantics of 3D shapes as they are progressively modeled within a pyramid architecture. These achievements in computer vision exemplify the efficacy of harnessing 3D point cloud data for gaining profound insights into spatial information.

Despite the successful application of pretraining on 3D point clouds in fields like computer vision, where it has revolutionized tasks such as object recognition, scene understanding, and autonomous navigation, its adoption in the domain of protein research remains limited. Our research bridges this gap by introducing an unsupervised pretraining method on 3D point clouds for proteins, aiming to leverage the rich spatial information encoded in these data to enhance protein representation learning. While 3D point clouds offer significant advantages in capturing the spatial structure of proteins, they face challenges in directly capturing other important features, such as residue types, sequence order, and residue-level interactions, which are essential for learning comprehensive and informative representations of proteins.

### E. Symmetry-preserving & Equivariant models

Symmetry in mathematics is a type of invariance: the property that a mathematical object remains unchanged under a set of operations or transformations. Such transformations can be either smooth, continuous, or discrete. Symmetries are important in many scientific problems and machine learning tasks. In graph representation learning, the scalar output of any graph neural networks must be invariant with respect to permutation of nodes [25, 38]. In chemistry and biochemistry, any neural networks predicting the molecular properties must be rotationally invariant with respect to the molecule’s orientation in space [39, 40]. In our work, our pretraining method on residue-level graphs respects the permutation symmetry while our pretraining method on 3D point cloud of the protein’ atoms respects both the rotation and translation symmetries.

### F. Multimodal representation learning

Protein representation learning is a challenging task because proteins are complex biomolecules with a hierarchical structure (as illustrated in 1). Each of these structural levels corresponds to a specific modality of representation, and learning on each individual modality has its strengths and limitations. While amino acid sequences offer insights into primary structural information, they may not capture the complex spatial relationships within protein structures. In contrast, residue-level graphs can provide crucial spatial information but may fall short in capturing long-range interactions in large protein structures. Similarly, 3D point clouds are excellent for capturing detailed spatial information, including the positions and arrangement of atoms, but they face challenges in directly capturing information about residues, such as residue types, sequence order, or residue-level interactions. To overcome these individual limitations and harness the strengths of different protein modalities, multi-modal learning has garnered attention. This approach holds the promise of generating comprehensive and enriched protein representations that can effectively address the multifaceted aspects of proteins and their functions.

Previous research has thoroughly explored the potential of Previous research extensively explored the potential of multimodal presentation learning to enhance our comprehension of proteins. For instance, the Protein Multimodal Network (PMN) architecture [15] was developed to unify various protein modalities, including sequences and 3D structures, aiming to improve the accuracy of predicting binding affinities. Additionally, a novel approach to structure-aware protein self-supervised learning [41] was introduced. This method entailed pretraining a graph neural network model to preserve protein structural information through self-supervised tasks, considering pairwise residue distances and dihedral angles. Self-supervised learning was further enhanced by leveraging a pretrained protein language model to establish a connection between sequential information in the language model and structural information in the graph neural network. Furthermore, the introduction of CoupleNet [42] presented a network explicitly designed to seamlessly integrate protein sequence and structure, effectively generating informative representations of proteins. This network proficiently combined residue identities and positions from sequences with geometric features derived from tertiary structures. Collectively, these innovations significantly contribute to our understanding of proteins and their diverse characteristics.

Previous works applying multimodal presentation learning to proteins have demonstrated remarkable effectiveness within the scope of the modalities they incorporated. However, there remains untapped potential in leveraging the full diversity of protein modalities, while also preserving the critical symmetries inherent to proteins, including permutation, rotation, and translation. In this study, we take a significant step forward by combining all three protein modalities, including amino acid sequences, residue-level graphs, and 3D point clouds, in a manner that preserves these symmetries. This approach allows us to create a truly comprehensive and multi-faceted protein representation that maximizes the unique strengths of each modality, which can then be used to improve the performance of a variety of protein-related tasks. We believe that our approach has the potential to make a significant contribution to the field of multimodal protein representation learning.

## III. METHOD

### A. Pretraining on Sequences

#### 1. Evolutionary Scale Modeling (ESM-2)

To obtain protein sequence representations, we employ the well-established pretrained Evolutionary Scale Modeling (ESM-2) [9]. ESM-2, a recent development building upon the ESM-1 architecture, is a state-of-the-art transformer architecture that offers variations with parameter counts ranging from 8 million to a staggering 15 billion. It is trained on over 65 million unique protein sequences to predict the identity of randomly masked amino acids. By leveraging a massive-scale training approach that involves solving missing puzzles, ESM-2 is able to effectively internalize complex sequence patterns across evolution and generate high-quality embeddings that are rich in both evolutionary and functional insights. Notably, the process of generating ESM-2 embeddings for a protein sequence is significantly more efficient in terms of computational resources and time investment as it does not rely on multiple sequence alignments (MSAs). ESM-2 embeddings are invaluable for various protein-related tasks, including structure prediction, design, and functional annotation, due to their computational efficiency and the profound evolutionary and functional insights they encapsulate.

#### 2. Our ESM-2 embeddings process

In our approach, it’s important to emphasize that we don’t engage in training the ESM-2 model ourselves. Instead, we make efficient use of several pre-trained ESM-2 checkpoints. To maintain a balanced model complexity across our modalities, we opted for a medium-sized ESM-2 version, which comprises 150 million parameters. The process initiates with the tokenization of protein sequences, facilitated by the corresponding tokenizer. Subsequently, we feed these tokenized sequences into the ESM-2 model to generate encoded representations. Specifically, we extract the last hidden state from the final model layer to obtain the encoded sequences, encapsulating vital information of proteins. This approach not only conserves computational resources but also ensures a consistent level of model complexity across our modalities.

### B. Pretraining on residue-level graphs

#### 1. Graph construction

To construct residue-level graphs, we employ a systematic approach to encapsulate the intricate spatial relationships between amino acid residues within protein structures. This process involves two key components:

- **Node Representation:** Within the residue-level graphs, each amino acid residue is treated as a node. These nodes are uniquely characterized through one-hot encoding, capturing the specific amino acid type for each residue. This representation provides a fundamental basis for the graph’s structural understanding.
- **Edge Construction:** To model intricate spatial dependencies, a K-nearest neighbor (KNN) edge construction strategy is applied. In this approach, we consider the alpha carbon atom coordinates as representative spatial points for each residue. By doing so, we can identify the k closest neighboring residues in 3-dimensional space, effectively linking residues within close spatial proximity. This strategy enables the graph to comprehensively model immediate structural interactions between amino acid residues.

#### 2. Variational Graph Autoencoder (VGAE)

In our pursuit of meaningful residue-level graph representations for protein representation learning, we utilize the Variational Graph Autoencoder (VGAE) model [10]. VGAE is a specialized framework for unsupervised learning on graph-structured data, building upon the principles of the variational autoencoder (VAE) [43].

This approach utilizes latent variables, empowering it to acquire interpretable latent representations tailored for undirected, unweighted graphs. Our VGAE architecture (as illustrated in Figure 3) consists of a graph convolutional network (GCN) encoder and a straightforward inner product decoder. The GCN encoder learns to encode the residue-level graph into a latent representation that captures the important structural features of the protein. The inner product decoder then reconstructs the residue-level graph from the latent representation. We pretrain our VGAE model to learn meaningful latent embeddings on a link prediction task on a dataset of residue-level graphs constructed from a set of protein structures.

**FIG. 2.**
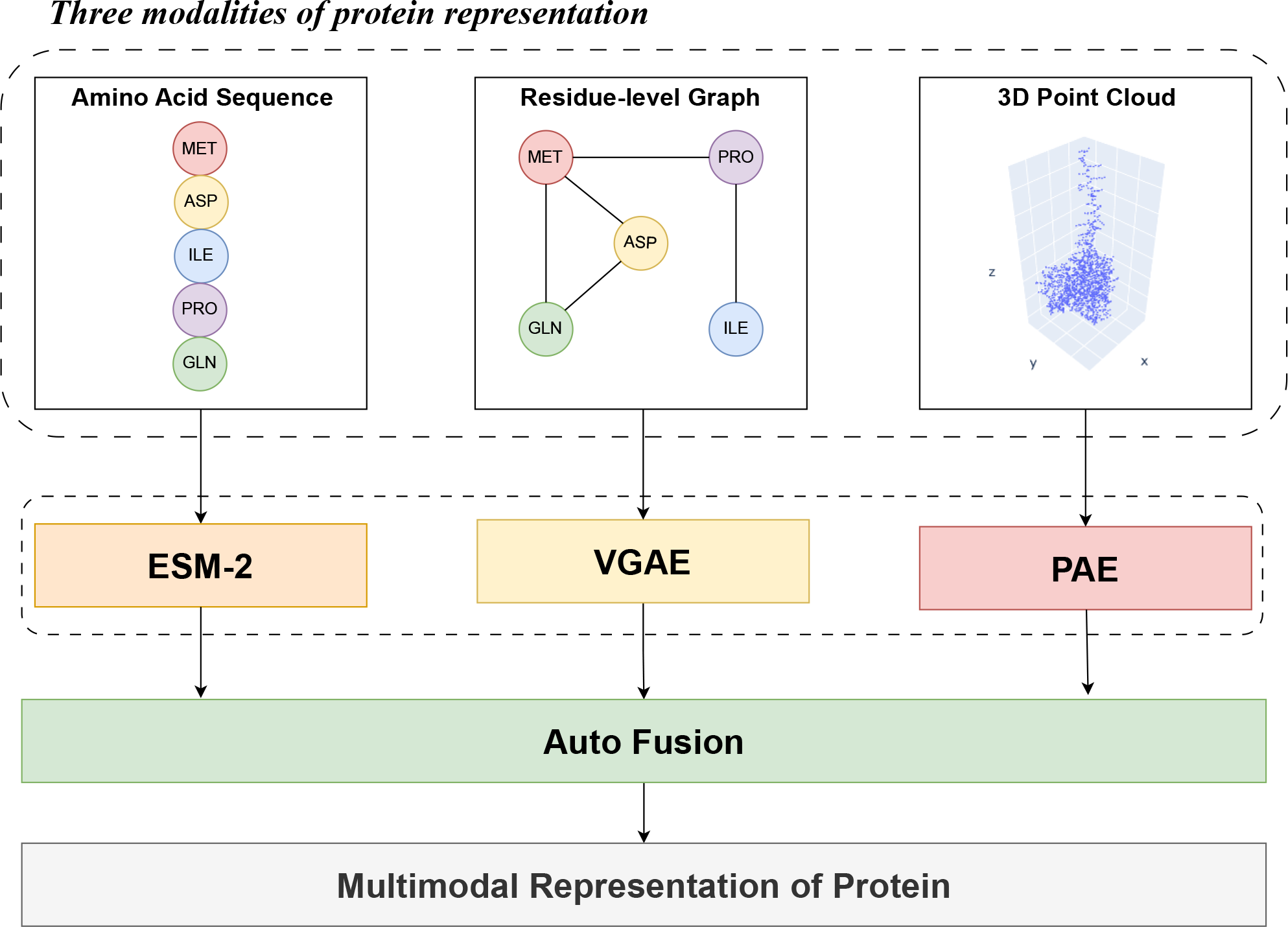
Overview of our multimodal protein representation learning framework. Our framework leverages symmetry-preserving multimodal unsupervised pretraining on proteins to learn informative representations that capture the intricate structural and functional features of proteins. To this end, we develop specialized pretraining methods for each protein modality, ensuring that the learned representations preserve the symmetry of the protein. We employ Evolutionary Scale Modeling (ESM-2) to capture essential information from amino acid sequences. To comprehensively understand the intricate spatial relationships between amino acid residues, we apply Variational Graph Auto-Encoders (VGAE) to residue-level graphs. Additionally, we utilize PointNet Auto-Encoder (PAE) to extract spatial arrangements of atoms from 3D point clouds. Once we have obtained meaningful representations from these diverse pretraining strategies, the next critical step in our framework is to fuse these representations using Auto-Fusion. This fusion process aims to synthesize joint representations from the pretrained models, capturing and combining the essential aspects of protein structures from multiple modalities. The fused multimodal representation can then be leveraged for a variety of protein-related downstream tasks.

**FIG. 3.**
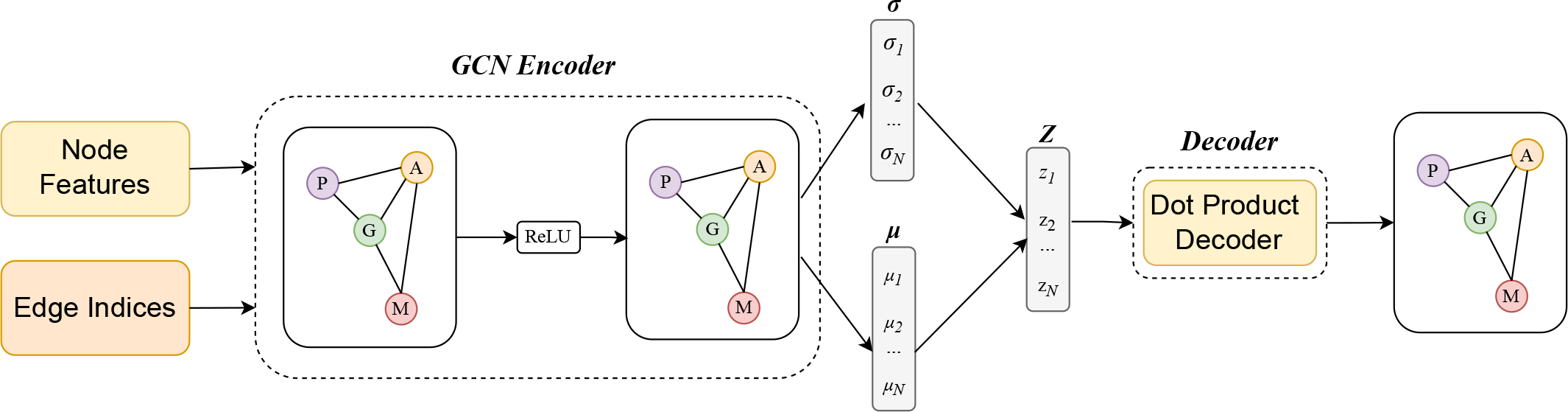
Architecture of the Variational Graph Autoencoder (VGAE) model. The input consists of edge indices and node features, composing a residue-level protein graph. The VGAE adopts a variational approach using the Graph Convolutional Network (GCN) Encoder, which is a key component of the model. In this encoder, GCN with two layers processes the input, applying ReLU activation, and produces both mean ***µ*** and log-variance **log(*σ*)** vectors. These vectors represent the parameters of the latent distribution. During training, latent vectors ***Z*** are sampled from this distribution, facilitating the learning of meaningful representations. The VGAE leverages this latent space to reconstruct the input graph through a dot product decoder.

The primary objective during training is to minimize a reconstruction loss, which assesses the VGAE model’s ability to faithfully reconstruct the original residue-level graph from the learned latent representation. This loss computation involves the consideration of both positive edges, as specified by the provided edge index, and negative edges, which are randomly generated through a negative sampling process. The reconstruction loss can be mathematically expressed as follows:

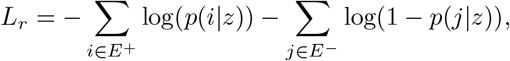

where E^+^ is the set of positive edges in the original residue-level graph, E^−^ is the set of negative edges randomly generated through negative sampling, and p(i|z) is the probability of edge i being present in the reconstructed residue-level graph, given the latent representation z. While traditional VAE-based models incorporate the Kullback-Leibler (KL) divergence loss to encourage the latent distribution to align with a standard normal distribution, we intentionally exclude this component. This strategic choice aligns with our precise focus on protein structure representation, allowing the VGAE to serve this purpose effectively.

Notably, our VGAE model preserves protein symmetry, including permutation, rotation, and translation. This preservation is achieved through a combination of design choices in the model. Firstly, the use of one-hot encoding for node features ensures that our model remains invariant to rotation and translation. One-hot encoding assigns a unique value to each amino acid type, making it insensitive to the orientation and position of residues within the protein structure. Secondly, the K-nearest neighbor (KNN) edge construction strategy, which relies on internode distances, maintains invariance to rotation and translation. This approach captures the spatial relationships between residues based on their relative distances, making the edge connections invariant to rotation and translation. Finally, Graph Convolutional Networks (GCNs) based on message passing scheme [28] is utilized to encode the residue-level graph into a set of latent vectors, each latent vector is associated with a residue / amino-acid, to ensure permutational equivariance. Furthermore, our graph decoder is implemented by taking the outer product of the latent matrix with itself (similar to VGAE in [10]), that ensures the whole auto-encoding processing is end-to-end permutation equivariant. These collective characteristics empower our model to effectively preserve the intricate symmetry within protein structures, allowing it to withstand various transformations and deliver precise protein representations.

### C. Pretraining on 3D point clouds

#### 1. Point clouds construction

Point cloud construction is a process of converting protein structural data into a point cloud format, which accurately captures the 3D spatial distribution of atoms within protein molecules. Our systematic construction process involves the following key components:

- Point Representation: Within our constructed point clouds, each atom from the protein structure is individually represented as a point in three-dimensional space, characterized by its specific coordinates (x, y, z). This detailed encoding captures the spatial identity of each atom and forms the fundamental basis for our point cloud representation, facilitating a comprehensive understanding of the structural intricacies within the protein.
- Standardization: By centering the coordinates around the origin and scaling the entire point cloud to fit within a unit sphere, we make the point clouds uniform in terms of their spatial distribution and size. This standardization is crucial for our framework as it allows for consistent and reliable analysis across different protein structures. It prevents variations in the spatial scales of point clouds, making them directly comparable and facilitating the learning process in subsequent stages of our model.
- Fixed number of points: Our point cloud representations are designed with a fixed number of points to maintain consistency during model training. When the number of atomic coordinates does not match the desired count, the construction procedure performs necessary actions, either padding the point cloud with zero vectors or, when fewer points are required, trimming the representation to the specified number.

#### 2. PointNet Autoencoder (PAE)

In our quest to derive informative 3D point cloud representations for protein representation learning, we adopt the PointNet Autoencoder (PAE) [11]. PAE, a specialized architecture designed for unsupervised learning using point cloud data, takes inspiration from the principles of the autoencoder (AE) while harnessing the robust capabilities of the PointNet framework [44]. Our PAE architecture (as illustrated in Figure 4) consists of two fundamental components: the PointNet encoder and decoder. The encoder’s primary role is to capture and extract essential features from the input 3D point cloud data. Within this component, PointNet assumes a pivotal role, enabling the model to comprehend and represent intricate structural information. On the other hand, the decoder takes the encoded representations and efficiently reconstructs the 3D point cloud data. It employs several layers, including fully connected layers and batch normalization, to ensure the accurate restoration of the point cloud’s spatial information.

**FIG. 4.**
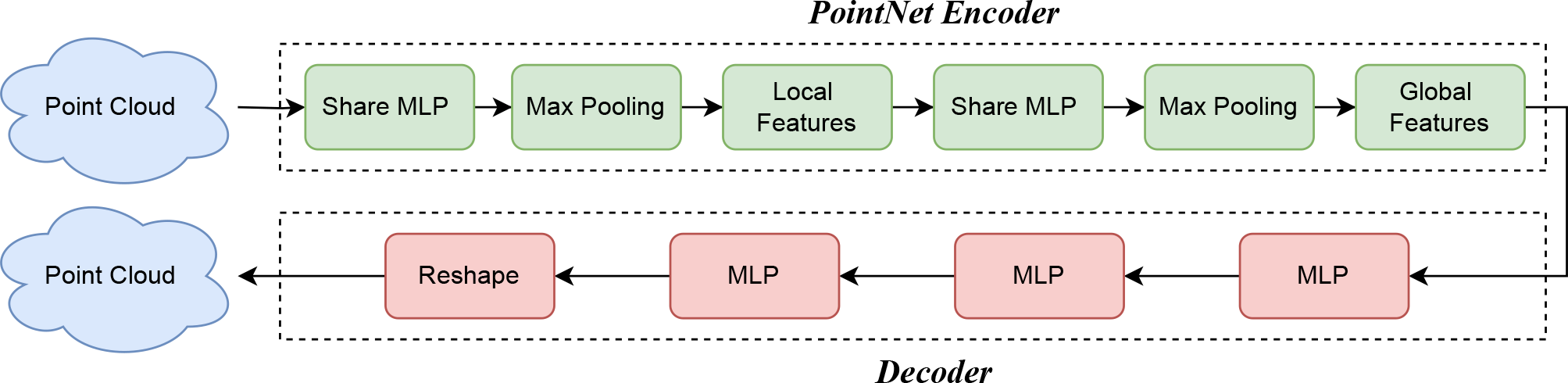
Architecture of the PointNet Autoencoder (PAE) model. The model consists of a PointNet-based encoder for capturing the structural information of the point cloud and a custom decoder for reconstructing the input point cloud from the encoded representation. The decoder is implemented as a multilayer perceptron (MLP) with hidden units of 512, 256, and 256 in its fully connected layers, each followed by ReLU activations and batch normalization. The final restoration is obtained by reshaping the output to match the specified number of points.

Our PAE were trained using the Chamfer Distance (CD) as the loss function. This metric is particularly effective for measuring the similarity between two point clouds and possesses the advantageous property of being invariant to the order of points within the clouds. The Chamfer Distance is calculated as the sum of squared distances between each point in one of the two point clouds and its nearest corresponding point in the other point cloud. This distance metric is expressed mathematically as follows:

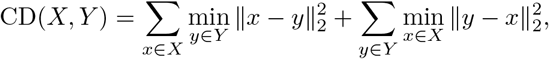

where X and Y are sets of points representing point clouds.

Our PAE model is noteworthy for maintaining protein symmetry, encompassing translation, rotation, and permutation. This preservation is accomplished via a combination of the model’s design decisions. Firstly, the PointNet architecture, which forms the foundation of our model, is a permutation-equivariant neural network architecture that can learn representations of point clouds that are invariant to the order of the points in the cloud. This is achieved through the use of a symmetric function in the max-pooling operation. Consequently, autoen-coders incorporating encoder stages inspired by Point-Net inherit the valuable capability to process unordered point clouds directly. This property proves to be an essential advantage, particularly in scenarios where point cloud order is not predefined. Secondly, the model benefits from random rotation during training, further enhancing its ability to capture and preserve the rotational symmetry of protein structures. Finally, the preprocessing step centers the coordinates of any input point cloud around the origin, eliminating concerns about translation symmetries, as the model will only be able to see the relative positions of the points in the point cloud, and not their absolute positions. These combined design choices ensure the comprehensive preservation of protein symmetry, making our PAE architecture highly effective in protein representation learning.

### D. Multimodal fusion

#### 1. Processing learned representations from pretrained models

A critical aspect of our approach is to harmonize the learned representations from various pretrained models to maintain balance across modalities. To achieve this, we ensure that all feature vectors obtained from different modalities have the same number of dimensionality. For the ESM and PAE models, their output is a feature vector consisting of 640 elements, but for the VGAE model, its output is a feature matrix with each row representing a feature vector for each node in the protein graph. To ensure uniform dimensionality, we employ the top-k pooling technique to select the top 640 nodes from the VGAE output matrix and compute an average feature vector. Furthermore, we standardize these feature vectors using z-score normalization to ensure that they are on the same scale and have the same distribution. This standardization improves the performance of the fusion model by preventing any single modality from dominating the fusion process.

#### 2. Auto-Fusion

In this study, we leverage Auto-Fusion [12] as our approach for multimodal synthesis, a method proven to enhance the model’s ability to extract intermodal features by optimizing the correlation between the different input modalities. Our Auto-Fusion architecture (as illustrated in Figure 5) consists of two primary components: the input feature fusion module and the reconstruction module. The fusion process commences by concatenating individual unimodal feature vectors, each obtained from our distinct pretraining methods. These concatenated features undergo a series of transformations facilitated by linear layers and non-linear activation functions within the input feature fusion module, culminating in the generation of an autofused latent vector. Subsequently, the reconstruction module takes over. Its primary objective is to reconstruct the original concatenated feature vector from the autofused latent representation. This process entails reverse transformations that seek to minimize the Euclidean distance between the original and reconstructed concatenated vectors using the Mean Squared Error (MSE), expressed as:

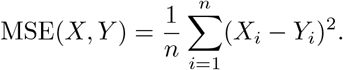

Notably, this reconstruction loss function plays a crucial role in refining the fused representation. Auto-Fusion ensures that the learned autofused vector retains only the essential shared information from the input modalities. It achieves this by effectively eliminating any arbitrary signals that might originate from the individual unimodal features. This meticulous optimization process enhances the quality of the fused representation, making it highly effective for synthesizing multimodal data. Our choice of Auto-Fusion aligns seamlessly with our goal of achieving a balanced and informative fusion of pretraining strategies in our framework.

**FIG. 5.**
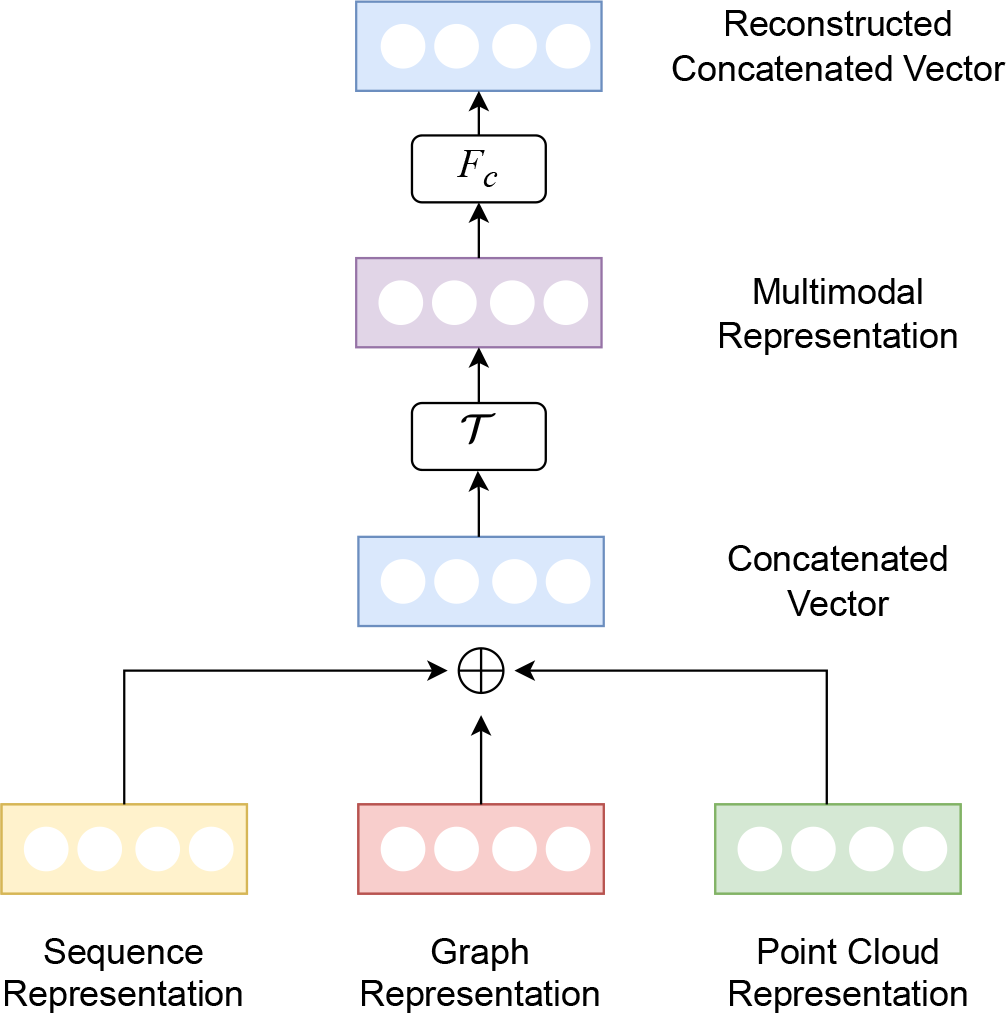
Architecture of the Auto-Fusion model. The sequence, graph, and point cloud representations are feature vectors generated by respective pretrained models for each modality. Initially, these vectors are concatenated and passed through a transformation ***T***, yielding a unified multimodal representation. The final reconstructed concatenated vector is obtained by processing this unified representation through another transformation, denoted as ***F***_***C***_.

## IV. EXPERIMENTS

### A. Pretraining evaluation

#### 1. Material

To facilitate robust unsupervised training, a substantial dataset comprising a multitude of PDB files is essential to ensure our model captures the intricate structure of proteins. In this endeavor, we utilized the Swiss-Prot structure dataset, which was sourced directly from the AlphaFold Protein Structure database. This dataset, generated by the AlphaFold platform, one of the state-of-the-art methods for protein folding, contains 542,378 PDB files, representing a diverse range of proteins from a variety of organisms. This makes it an ideal dataset for unsupervised pretraining, as it allows the model to learn a generalizable representation of protein structures. To organize this vast dataset effectively, we divided it into subsets, comprising a training set, a validation set, and a test set. This division follows a balanced ratio of 70:20:10, enabling us to train, validate, and evaluate our model on distinct portions of the dataset and ensure its robustness and generalization. For each of the subsequent pretraining steps, including VGAE, PAE, and Auto-Fusion, we maintained a consistent pretraining duration of 100 epochs. This standardized approach ensures that all models had an equal opportunity to learn informative protein representations during the pretraining phase.

#### 2. Pretraining evaluation on VGAE

During the training process, both the training loss and validation loss consistently decreased, indicating effective learning. The final training loss and validation loss for VGAE converged to 0.951 and 0.952, respectively. For performance evaluation, VGAE was assessed on the link prediction task using the test dataset. The results indicate a remarkable achievement, with an Area Under the Curve (AUC) of 0.95 and an Average Precision (AP) of 0.97. These metrics emphasize VGAE’s effectiveness in capturing essential structural features in protein graphs and demonstrate its ability to predict protein interactions accurately.

#### 3. Pretraining evaluation on PAE

Throughout the training process, the training loss exhibited a consistent and significant decrease, demonstrating the model’s ability to effectively comprehend intricate protein structures. However, the validation loss curve displayed some instability during this phase, albeit with an overall trend towards decreasing values compared to the initial levels. The final training loss and validation loss for PAE were 6.51 and 7.84, respectively. The relatively high variance observed in the validation loss curve could potentially be attributed to several factors, such as the complexity and diversity within the protein structure dataset. The PAE model achieved a Chamfer distance of 7.89 on the test set for the task of reconstructing the 3D point cloud of proteins. This is an outstanding result that demonstrates the ability of the PAE model to learn informative representations of protein structures, even for proteins with complex and diverse structures.

#### 4. Pretraining evaluation on Auto-Fusion

During the training process of Auto-Fusion, the training and validation loss consistently decreased over time, albeit with occasional fluctuations. The final training loss and validation loss for Auto-Fusion converged to and 0.299, respectively. This behavior indicates the model’s adaptability to complex protein structures while maintaining a general decreasing trend in loss values. Auto-Fusion’s performance was further assessed using a dedicated test dataset, with a focus on its capability to reconstruct the original concatenated feature vector from the autofused latent representation. Remarkably, Auto-Fusion achieved an exceptional Mean Squared Error (MSE) of 0.03. This low MSE value emphasizes the model’s efficacy in preserving essential structural features during the reconstruction process, solidifying Auto-Fusion’s role as a robust component in our framework for protein representation learning.

### B. Fine-tuning results

#### 1. Protein-ligand binding affinity prediction

##### a. Problem statement

Structure-based drug design (SBDD), a powerful approach for identifying potential drug candidates, involves assessing the fit and interactions of small molecules (ligands) within the binding sites of target proteins [45]. The strength of these interactions, known as binding affinity, is a key determinant of a ligand’s ability to modulate the protein’s biological function [46]. Therefore, compounds with high binding affinity to target proteins are prioritized as potential drug candidates. The accurate prediction of binding affinity is essential for efficiently screening compound libraries and optimizing lead compounds, thereby reducing the costs of drug discovery.

##### b. Material

To gauge the effectiveness of our multi-modal representation in predicting protein-ligand binding affinity, we conducted assessments using three distinct ligand-binding datasets: DAVIS [47], KIBA [48], and PDBbind v2020 [49]. The DAVIS dataset encompasses 442 proteins and 68 ligands, forming a dataset comprising 30,056 protein-ligand binding pairs. In DAVIS, the binding scores are quantified as KD constants. Conversely, the KIBA dataset is characterized by 229 proteins and an expanded collection of 2,111 ligands, creating a dataset that consists of 118,254 protein-ligand binding pairs, with binding affinities represented as KIBA scores. The PDBbind v2020 dataset extends our evaluation with a rich repository of 19,433 protein-ligand binding pairs, divided into a general set with 14,127 samples and a refined set with 5,316 samples. This dataset provides experimentally measured binding affinities for protein-ligand interactions (PLI), encompassing data on peptides and nucleic acids expressed in units like Kd, Ki, or IC50, or their negative logarithmic equivalents (Pkd). The incorporation of these diverse datasets forms a comprehensive and varied tested, allowing us to thoroughly assess the predictive capabilities of our multimodal representation when estimating protein-ligand binding affinities. To ensure a fair and standardized evaluation, we meticulously followed the test/training/validation split settings as outlined in previous studies, specifically adhering to the configurations defined in the respective sources for the DAVIS, KIBA, and PDBbind v2020 datasets [15, 50]. By maintaining this consistency, we aimed to create a level playing field for comparisons, allowing for an equitable assessment of our multimodal representation’s performance. Furthermore, we adopted the same measurement metrics used in these previous works, aligning our methodology with established standards to facilitate straightforward and meaningful comparisons with existing research outcomes. This rigorous approach enhances the reliability and comprehensibility of our results within the broader scientific community.

##### c. Method

For each data point within these datasets, we used a hybrid approach that combined the protein representation from our framework with the ligand representation from Morgan fingerprints. We concatenated these two distinct representations into a single feature vector, which then used as input to our Gaussian Process Regressor model. The Gaussian Process Regressor is a powerful machine learning model that leverages the principles of Gaussian processes for regression tasks. It is particularly well-suited for modeling complex relationships between input features and target values. By employing GPR as our modeling technique, we harnessed its ability to capture intricate patterns within the data, enabling us to make accurate predictions while quantifying the associated uncertainty. This approach allowed us to effectively estimate protein-ligand binding affinities and assess ligand-protein interactions within the given datasets.

#### 2. Protein fold classification

##### a. Problem statement

Protein fold classification plays a crucial role in unraveling the profound interplay between protein structure, function, and the evolutionary trajectory of biological molecules. It enables the grouping of proteins into specific fold classes, drawing upon common attributes such as their secondary structure composition, spatial arrangements, and connectivity patterns. This classification process is indispensable for the comprehensive understanding of proteins. By categorizing them into fold classes, we gain insights into their functional characteristics, allowing us to decipher the intricate relationship between form and function. Additionally, it provides valuable information about how proteins have evolved over time. In light of these considerations, our primary objective in this context is to accurately predict the fold class to which a given protein belongs.

##### b. Material

For protein fold classification task, we utilized the SCOPe 1.75 dataset, as established by [76], which offers well-defined training, validation, and test partitions. This dataset encompasses a comprehensive collection of 16,712 proteins originating from 1,195 unique protein folds. The 3D structural information for these proteins was sourced from the SCOPe 1.75 database, as provided by [77]. The dataset comprises three distinct test subsets: ‘Fold’ where proteins from the same superfamily are excluded from the training set; ‘Superfamily’ in which proteins from the same family are omitted from the training data; and ‘Family’ wherein proteins from the same family are retained within the training set.

##### c. Method

For each data point within this dataset, we exclusively leveraged the multimodal representation of proteins generated by our framework. This representation served as the sole input for our XGBoost Classifier model. XGBoost Classifier is a powerful machine learning algorithm that is based on the gradient boosting principle. Gradient boosting is an ensemble learning technique that combines the predictions of multiple weak learners to produce a more accurate prediction. By utilizing the XGBoost Classifier as our chosen model, we harnessed its strength in making accurate and robust predictions, even when dealing with intricate protein structural data. This approach allowed us to effectively classify proteins into their respective fold classes based on the provided multimodal representations. To assess the performance of our model in protein structural classification, we employed the accuracy metric.

#### 3. Enzyme identification

##### a. Problem Statement

There are seven primary protein categories that encompass all proteins, each serving a unique function in biological processes. These categories consist of antibodies, contractile proteins, enzymes, hormonal proteins, structural proteins, storage proteins, and transport proteins. Among these diverse protein types, enzymes hold a pivotal position. Enzymes are proteins whose function is to catalyze, or accelerate, specific chemical reactions within the cell. The accurate identification of enzymes within the larger spectrum of proteins is fundamental to understanding the intricate biochemical workings of life and has significant implications for various fields, such as biotechnology and medicine.

##### b. Material

In this task, we utilized the D&D benchmark dataset defined by [85], which consists of 1178 structurally diverse proteins, comprising 691 enzymes and 487 non-enzymes, based on annotations in the PDB and Medline abstracts. To ensure a fair and consistent comparison with prior research, we adopted the 10-fold cross-validation partitioning established in [80].

##### c. Method

Similar to our approach to protein fold classification, we utilized our multimodal representation learning framework to extract informative protein representations for the enzyme identification task. These representations were then fed into an XGBoost Classifier model to predict whether a given protein is an enzyme or not. Our approach allowed us to effectively classify proteins into their respective categories, distinguishing between enzymes and non-enzymes based on the provided multimodal representations. To measure the performance of our model in this task, we calculated the average accuracy across the 10-fold cross-validation setup.

#### 4. Mutation stability prediction

##### a. Problem statement

Predicting the stability of protein mutations is a critical task in understanding the intricate interplay between genetic variations and protein structure and function. Mutations can profoundly alter protein stability, leading to changes in their structural dynamics and, consequently, affecting their biological activities. Accurately predicting the stability of mutated proteins is essential for designing novel proteins. While experimental techniques for probing these changes are labor-intensive, the development of efficient computational methods presents a promising alternative. To address this challenge, the task is formulated as a binary classification problem, aiming to predict whether the stability of the protein complex increases or decreases as a result of the mutation.

##### b. Material

The dataset for the Mutation stability prediction (MSP) task is sourced from Atom3D [92], a comprehensive benchmark encompassing both novel and established datasets spanning several critical classes of biomolecules. Each mutation in the MSP task includes a PDB file with the residue of interest transformed to the specified mutant amino acid, as well as the native PDB file. A total of 4,148 mutant structures accompanied by their 316 wild-type (WT) structures are provided. For labeling, a value of 1 is assigned to a mutant if the dissociation constant (Kd) of the mutant protein is less than that of the wild-type protein, indicating improved binding; otherwise, a label of 0 is assigned. To ensure that the model is not simply memorizing the training data, protein complexes are such that no protein in the test dataset has more than 30% sequence identity with any protein in the training dataset.

##### c. Method

To tackle the mutation stability prediction task, we employed our multimodal representation learning framework to extract informative representations of both wild-type and mutant proteins. These representations encapsulate the structural and functional characteristics of the proteins, providing the necessary information to predict the stability changes induced by mutations. Subsequently, the extracted multimodal representations were concatenated to form a single, combined representation. This concatenation process enabled the model to integrate information from both wild-type and mutant proteins, allowing it to learn the relationships between their representations and predict mutation stability outcomes. The concatenated multimodal representation was then fed into an XGBoost Classifier model. The choice of the AUROC as the evaluation metric stems from its suitability for imbalanced datasets, ensuring robust performance assessment in the context of the inherent data imbalance.

## C. Discussion

The results of our experiment underscore the effectiveness of our multimodal representation learning framework for acquiring informative protein representations. By learning these representations through unsupervised pre-training models, our approach, while not surpassing state-of-the-art (SOTA) models, consistently delivers impressive results across 4 crucial downstream tasks: protein-ligand binding affinity prediction (see Table I and II), protein fold classification (see Table III), enzyme identification (see Table IV) and mutation stability prediction (see Table V). This highlights the versatility and potential of our approach for a wide range of protein-related tasks, making it a valuable resource for various applications within the field of protein research. Particularly noteworthy is the fact that we exclusively employ traditional machine learning methods like Gaussian Process and XGBoost to utilize our multimodal protein representation for these tasks, and yet, we consistently achieve highly favorable results. It demonstrates that our protein representations are general and informative enough to enable traditional ML models to achieve remarkable results on complex protein-related tasks, without the need for complex deep learning architectures.

**TABLE I.** Experimental results of Protein-ligand binding affinity prediction task on DAVIS and KIBA dataset.

| Approach | DAVIS |  |  | KIBA |  |  |
| --- | --- | --- | --- | --- | --- | --- |
| | MSE ↓ | CI ↑ | $r_m^2$ ↑ | MSE ↓ | CI ↑ | $r_m^2$ ↑ |
| KronRLS [51] | 0.379 | 0.871 | 0.407 | 0.411 | 0.782 | 0.342 |
| SimBoost [52] | 0.282 | 0.872 | 0.640 | 0.220 | 0.836 | 0.629 |
| SmCNN-DTA [53] | 0.319 | 0.852 | 0.590 | 0.274 | 0.821 | 0.573 |
| DeepDTAY [54] | 0.261 | 0.878 | 0.630 | 0.194 | 0.863 | 0.673 |
| WideDTA [55] | 0.886 | 0.202 | - | 0.875 | 0.179 | - |
| AttentionDTA [56] | 0.216 | 0.893 | 0.670 | 0.155 | 0.882 | 0.755 |
| MATT-DTI [57] | 0.227 | 0.891 | 0.653 | **0.150** | 0.882 | 0.756 |
| GraphDTA [58] | 0.258 | 0.884 | 0.656 | 0.162 | 0.879 | 0.736 |
| FusionDTA [59] | 0.220 | 0.903 | 0.666 | 0.167 | **0.891** | 0.699 |
| BiCompDTA [60] | 0.237 | 0.904 | 0.696 | 0.167 | **0.891** | 0.757 |
| PMN [15] | **0.202** | **0.906** | **0.739** | 0.153 | 0.874 | **0.767** |
| ESM (ours) | 0.300 | 0.690 | 0.311 | 0.258 | 0.789 | 0.326 |
| VGAE (ours) | 0.508 | 0.631 | 0.032 | 0.344 | 0.752 | 0.250 |
| PAE (ours) | 0.558 | 0.636 | 0.042 | 0.329 | 0.749 | 0.252 |
| Multimodal (ours) | 0.248 | 0.699 | 0.414 | 0.199 | 0.812 | 0.471 |

**TABLE II.**
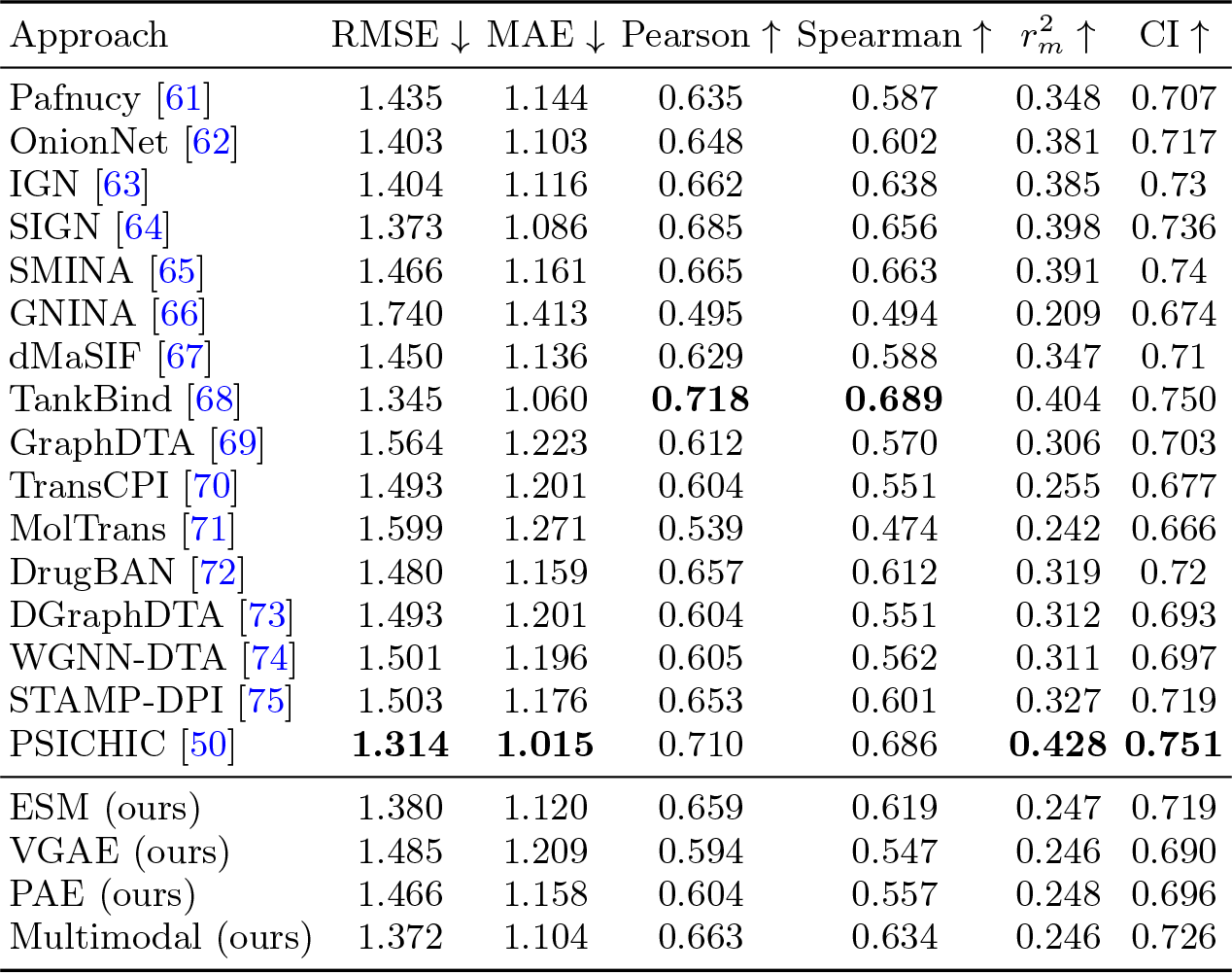
Experimental results of Protein-ligand binding affinity prediction task on PDBBind v2020 dataset.

**TABLE III.**
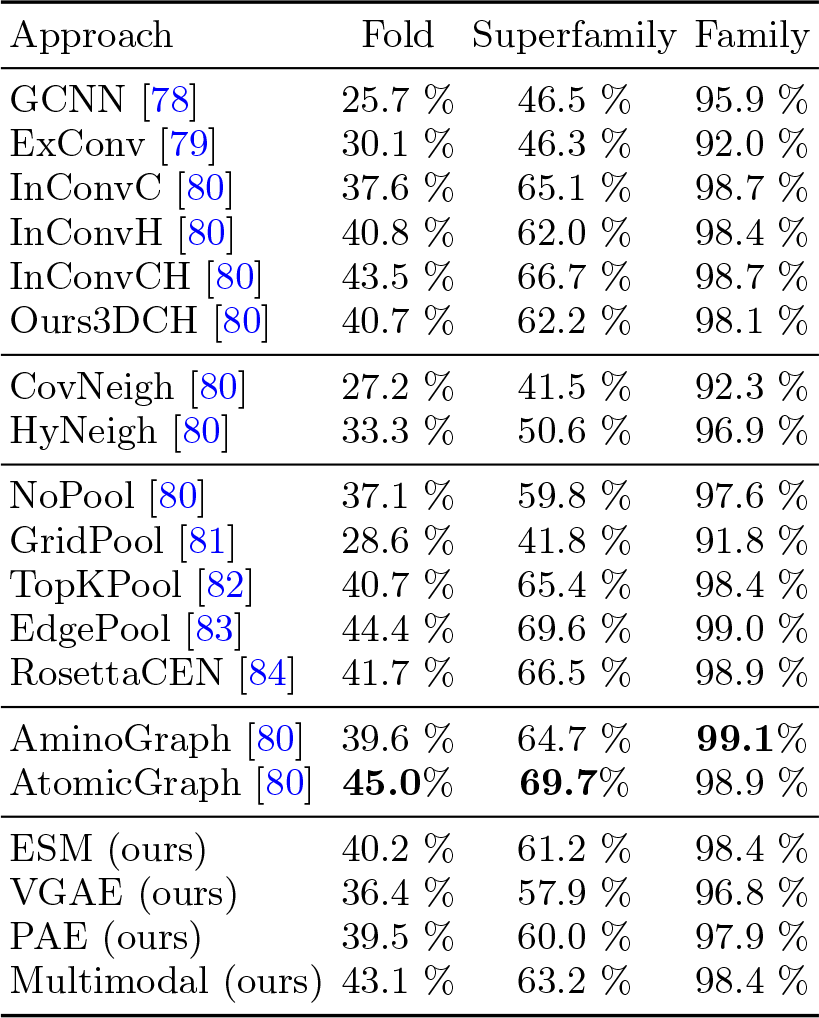
Experimental results of Protein fold classification task on SCOPe 1.75 dataset, reported in terms of Accuracy. Results are presented for three distinct test subsets: Fold, Superfamily, and Family.

| Approach | Fold | Superfamily | Family |
| --- | --- | --- | --- |
| GCNN [78] | 25.7 % | 46.5 % | 95.9 % |
| ExConv [79] | 30.1 % | 46.3 % | 92.0 % |
| InConvC [80] | 37.6 % | 65.1 % | 98.7 % |
| InConvH [80] | 40.8 % | 62.0 % | 98.4 % |
| InConvCH [80] | 43.5 % | 66.7 % | 98.7 % |
| Ours3DCH [80] | 40.7 % | 62.2 % | 98.1 % |
| CovNeigh [80] | 27.2 % | 41.5 % | 92.3 % |
| HyNeigh [80] | 33.3 % | 50.6 % | 96.9 % |
| NoPool [80] | 37.1 % | 59.8 % | 97.6 % |
| GridPool [81] | 28.6 % | 41.8 % | 91.8 % |
| TopKPool [82] | 40.7 % | 65.4 % | 98.4 % |
| EdgePool [83] | 44.4 % | 69.6 % | 99.0 % |
| RosettaCEN [84] | 41.7 % | 66.5 % | 98.9 % |
| AminoGraph [80] | 39.6 % | 64.7 % | **99.1%** |
| AtomicGraph [80] | **45.0%** | **69.7%** | 98.9 % |
| ESM (ours) | 40.2 % | 61.2 % | 98.4 % |
| VGAE (ours) | 36.4 % | 57.9 % | 96.8 % |
| PAE (ours) | 39.5 % | 60.0 % | 97.9 % |
| Multimodal (ours) | 43.1 % | 63.2 % | 98.4 % |

**TABLE IV.** Experimental results of Enzyme identification task on D&D dataset.

| Approach | Accuracy ↑ |
| --- | --- |
| Gao & Ji (2019) [86] | 84.4% |
| Ying et al. (2018) [87] | 82.1% |
| Zhao & Wang (2019) [88] | 82.0% |
| Nguyen et al. (2020) [89] | 81.2% |
| Zhang et al. (2019) [90] | 79.7% |
| Togninalli et al. (2019) [91] | 79.4% |
| Hermosilla et al. (2021) [80] | **85.5%** |
| ESM (ours) | 82.7% |
| VGAE (ours) | 81.4% |
| PAE (ours) | 80.4% |
| Multimodal (ours) | 83.9% |

**TABLE V.** Experimental results of Mutation stability prediction task on Atom3D.

| Approach | AUROC ↑ |
| --- | --- |
| 3DCNN [92] | 0.574 |
| GNN [92] | 0.609 |
| ENN [92] | 0.574 |
| Rao et al. (2019) [93] | 0.554 |
| ESM (ours) | 0.509 |
| VGAE (ours) | 0.463 |
| PAE (ours) | 0.488 |
| Multimodal (ours) | **0.612** |

As evidenced by our ablation study, our models follow a distinct order in their performance. The multimodal approach stands out as the most efficient in delivering consistently competitive results across all tasks. This efficiency arises from the fact that it benefits from the combined strength of different modalities, allowing it to capture a broader spectrum of protein features and, consequently, achieve robust and informative representations. ESM closely follows in terms of efficiency, maintaining a high level of performance. ESM’s efficiency is due to its ability to leverage pre-trained contextual embeddings, which capture the rich and intricate language of proteins. PAE and VGAE, while effective, exhibit complementary strengths and weaknesses, with PAE performing better on some specific tasks, while VGAE performs better on others. For example, PAE achieves better results on the protein-ligand binding affinity prediction task, protein fold classification and mutation stability prediction, while VGAE achieves better results on the enzyme identification task. These findings emphasize that for models that learned from a single modal, their effectiveness may be task-dependent. In contrast, our multimodal framework can achieve stable results across multiple tasks, reducing sensitivity to task-specific traits.

## V. CONCLUSION

In this paper, we have presented a comprehensive framework for protein representation learning, addressing critical challenges in the field. Our contributions encompass the development of dedicated unsupervised symmetry-preserving pretraining methods for distinct protein modalities, utilizing Evolutionary Scale Modeling (ESM-2) for amino acid sequences, Variational Graph Auto-Encoders (VGAE) for residue-level graphs, and PointNet Auto-Encoder (PAE) for 3D point clouds. Leveraging Auto-Fusion, our approach synthesizes joint representations, facilitating effective intermodal information extraction. Our experimental results demonstrate the effectiveness of multimodal unsupervised symmetry-preserving pre-training methods for learning protein representations, which is evidenced by our framework’s impressive performance on several protein-related downstream tasks, including protein-ligand binding affinity prediction, protein fold classification, enzyme identification and mutation stability prediction. Our framework’s ability to offer informative protein representations presents exciting opportunities for researchers to tackle complex problems in the realm of protein science.

## Appendix A: Multimodal Embeddings Visualization

In this appendix, we present t-SNE [94] visualizations of the multimodal embeddings encoded by our framework (Figures 6 and 7). These visualizations provide valuable insights into the effectiveness of our approach on two classification tasks: Protein fold classification and Enzyme Identification. While it’s crucial to acknowledge that the unsupervised nature of our embedding method may not always result in perfect class separation with all data points perfectly clustered together, it does reveal intriguing clustering patterns that suggest our capability to group a significant portion of data points belonging to the same class together effectively. These visualizations offer valuable insights into the distribution of our learned embeddings and highlight the substantial potential of our approach in capturing meaningful relationships among data points.

**FIG. 6.**
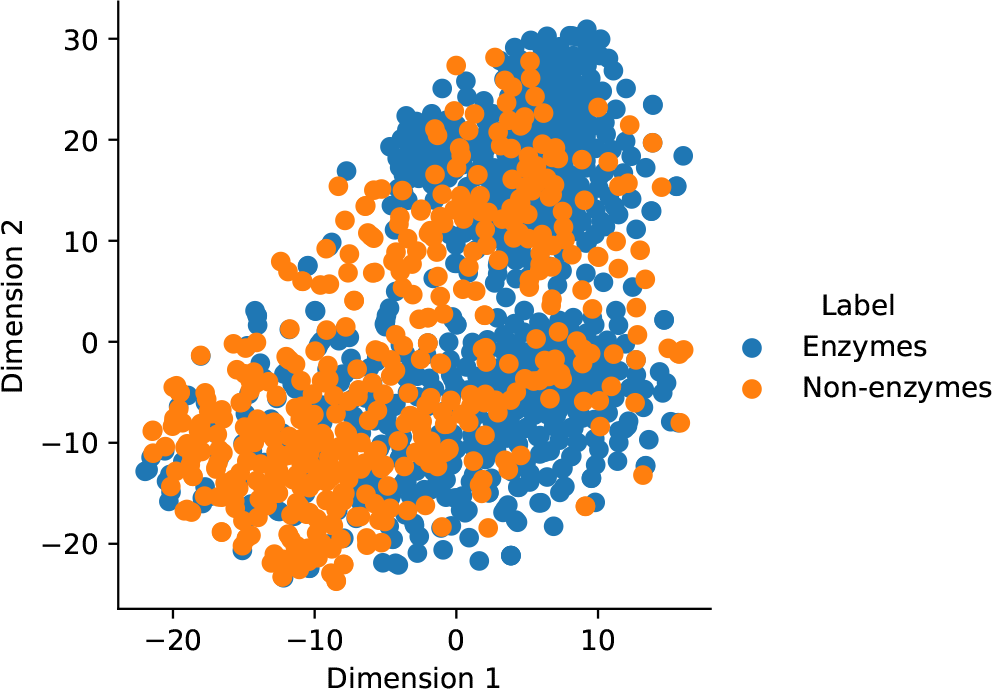
Visualization of the learned representations on the Enzyme Identification task (D&D dataset). Each point represents a protein, and the color indicates the protein’s enzyme class.

**FIG. 7.**
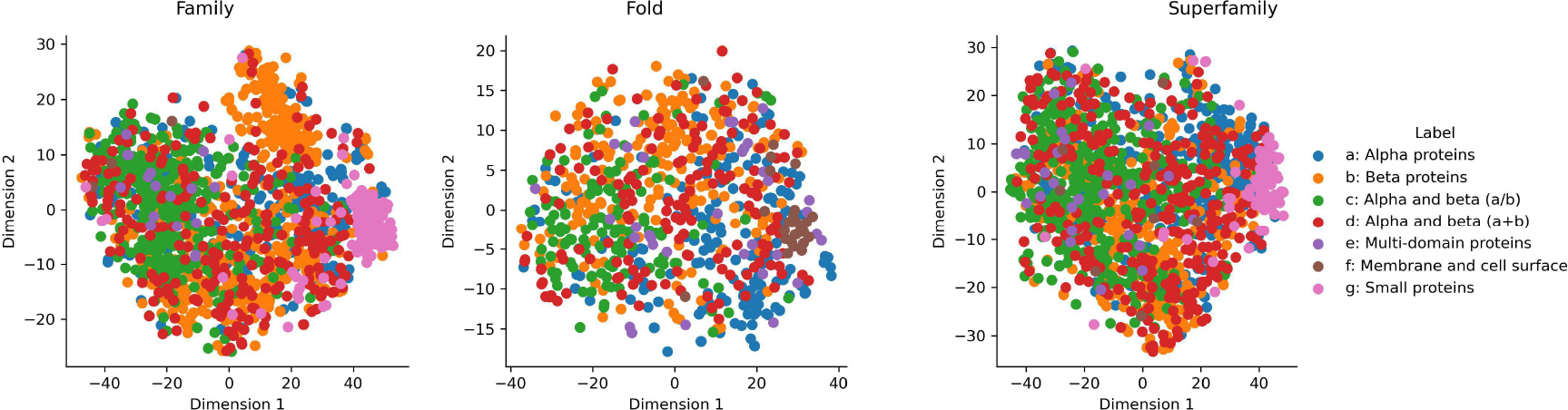
Visualization of the learned representations on the Protein Fold Classification task (SCOPe 1.75 dataset). Each point represents a protein, and the color indicates the protein’s fold classes.

## Appendix B: Implementation Details

This appendix provides an overview of the implementation details, with a focus on the key hyperparameters used in the models and the training process for our study. Understanding these details is essential for replicating and comprehending our experiments.

### 1. Pretraining Process

For the pretraining phase, we leveraged the ESM pretrained model, specifically opting for the 150 million parameter version available through Hugging Face’s Transformers [95] library. Our implementation of the VGAE, PAE, and Auto-Fusion models leveraged the PyTorch [96] and PyTorch Geometric [97] deep learning frameworks. Each pretrained model was configured to encode proteins into feature vectors, each comprising 640 elements. Subsequently, these individual feature vectors generated by the three models were synthesized into a unified feature vector with 1024 elements by our Auto-Fusion model. In terms of model training parameters, we conducted training over 100 epochs with a batch size of 256, employing a fixed learning rate of 0.001 and the Adam optimization algorithm. To enhance the training process’s effectiveness, we ensured random shuffling of the training data before initiating the training process.

### 2. Downstream Tasks

For our downstream tasks, we employed two well-established machine learning algorithms: Gaussian Process (GP) and XGBoost. The GP model found its application in protein-ligand binding affinity prediction, while the XGBoost model was utilized for both protein fold classification and enzyme identification. The GP model was implemented using the GPyTorch [98] library, a robust tool for constructing and training GP models within the PyTorch framework. The GP model employed the Gaussian likelihood, a fitting choice for regression tasks, and the Radial Basis Function (RBF) kernel. The selection of the RBF kernel was motivated by its ability to capture smooth and continuous variations in data, aligning well with the typical characteristics of protein-ligand binding affinities. The training process for the GP model aimed at optimizing model parameters to effectively fit the training data. To achieve this, a negative log-likelihood loss function was employed, and the GP model was trained for 100 epochs, utilizing a batch size of 256 and the Adam optimization algorithm. This iterative training process enabled the model to learn underlying data patterns and make accurate predictions for unseen protein-ligand binding affinities. In parallel, we configured the XGBoost [99] model with specific hyperparameters, including a learning rate of 0.1, a maximum tree depth of 3, and a total of 1000 estimators. During the training process, we integrated evaluation metrics, with multi-class logarithmic loss serving as the chosen evaluation metric to monitor the model’s performance. To prevent overfitting, early stopping was implemented, with a tolerance of 10 rounds. This configuration ensured the XGBoost model’s effectiveness in the tasks of classifying protein folds and identifying enzymes.

